# Pathogenic LRRK2 control of primary cilia and Hedgehog signaling in neurons and astrocytes of mouse brain

**DOI:** 10.1101/2021.03.02.433576

**Authors:** Shahzad S. Khan, Yuriko Sobu, Herschel S. Dhekne, Francesca Tonelli, Kerryn Berndsen, Dario R. Alessi, Suzanne R. Pfeffer

## Abstract

Previously, we showed that cholinergic interneurons of the dorsal striatum lose cilia in mice harboring the Parkinson’s disease associated, kinase activating, R1441C LRRK2 mutation (Dhekne et al., 2018). Here we show that this phenotype is also seen in two mouse strains carrying the most common human G2019S LRRK2 mutation. Heterozygous loss of the PPM1H phosphatase that is specific for LRRK2-phosphorylated Rab GTPases (Berndsen et al., 2019) yields the same cilia loss phenotype, strongly supporting a connection between Rab GTPase phosphorylation and cilia loss. In addition, astrocytes throughout the striatum show a ciliation defect in LRRK2 and PPM1H^-/+^ mutant models. Hedgehog signaling requires cilia, and loss of cilia correlates here with a loss in induction of Hedgehog signaling as monitored by in situ hybridization of *Gli1* transcripts. These data support a model in which LRRK2 and PPM1H mutant mice struggle to receive and respond to critical Hedgehog signals in the nigral-striatal pathway.

## Introduction

Mutations in the kinase encoded by the LRRK2 gene represent the predominant cause of familial Parkinson’s disease (PD), a neurodegenerative disorder that results in the loss of dopaminergic neurons in the substantia nigra pars compacta (Poewe et al. 2017; Alessi and Sammler, 2018). LRRK2 encodes a protein kinase and recent work has shown that a subset of Rab GTPases comprise its primary substrates (Steger et al., 2016; 2017). Pathogenic mutations localize to LRRK2’s ROC domain (eg. R1441C) and kinase domain (eg. G2019S), increase its kinase activity (West et al., 2005; Greggio et al., 2006; Jaleel *et al*., 2007; Ito *et al*., 2016; Steger *et al*., 2016), and interactions with other proteins including Rab29 (Kuwahara et al., 2016; Purlyte et al., 2018; Liu et al., 2018; Gomez et al., 2019) and VPS35 can also activate LRRK2 (Linhart et al., 2014; Mir et al., 2018). Reversal of LRRK2 phosphorylation is mediated at least in part by the phosphatase PPM1H that was recently discovered to specifically reverse LRRK2 action on multiple Rab GTPases (Berndsen et al., 2019). In cell culture, loss of PPM1H in wild type mouse embryonic fibroblast (MEF) cells phenocopies the loss of cilia seen upon expression of pathogenic LRRK2 (Berndsen et al., 2019).

Rab GTPases are master regulators of protein trafficking and carry out their roles by binding to specific partner proteins when the Rabs are GTP-bound (Pfeffer, 2017, 2018). Phosphorylation of Rab proteins interferes with their abilities to be loaded with GTP by cognate guanine nucleotide exchange factors, a prerequisite for their binding to partner effector proteins (Steger et al., 2016, 2017). This alone would interfere with normal Rab GTPase function. Strikingly, once phosphorylated, Rab GTPases switch their preference and bind to new sets of phospho-specific protein effectors. For Rab8 and Rab10 these include RILPL1, RILPL2, JIP3 and JIP4 proteins (Steger et al., 2017; Dhekne et al., 2018; Waschbüsch et al., 2020) and Myosin Va (Dhekne et al., 2021). The consequences of new phospho-Rab interactions include pathways by which LRRK2 blocks ciliogenesis in cell culture and mouse brain via a process that requires RILPL1 and Rab10 proteins (Steger et al., 2017; Dhekne et al., 2018); centriolar cohesion is also altered (Madero-Pérez et al., 2018; Ordonez et al., 2019).

Although later stages of PD resemble those seen in Alzheimer’s disease, PD is first and foremost a movement disorder, and studies to understand it’s underlying causes must focus on understanding why PD is specifically characterized by dopaminergic neuron loss in the substantia nigra. LRRK2 is most highly expressed in immune cells, lung, kidney and intestine, but it is also present in varying levels throughout the brain (Lis et al., 2018; West et al., 2014; Mandemakers et al., 2012). The striatum is comprised primarily of medium spiny neurons, interneurons and glial cells such as astrocytes. In an important study, Gonzalez-Reyes et al. (2012) showed that dopaminergic neurons in the substantia nigra secrete Sonic Hedgehog (Hh) that is sensed by poorly abundant, cholinergic interneurons in the striatum. Hh is needed for the survival of both these cholinergic target cells and the Hh-producing dopaminergic neurons, despite the fact that only the cholinergic neurons express the PTCH1 Hh receptor. Gonzalez-Reyes et al. (2012) showed further that Hh triggers secretion of glial derived neurotrophic factor (GDNF) from the cholinergic neurons, which provides reciprocal neuroprotection for the dopaminergic neurons of the substantia nigra.

We found previously that the rare, striatal, cholinergic interneurons that would normally sense Hh via their primary cilia are less ciliated in mice carrying the R1441C LRRK2 mutation (Dhekne et al., 2018). In that study, we proposed that cilia loss would decrease the ability of these cells to sense Hh signals. We show here that Hh signaling is indeed impacted by cilia loss in multiple LRRK2 mutant mouse models, even as early as 10 weeks of age. Moreover, we show that striatal astrocytes share a broad ciliary deficit that likely impacts synaptic function.

## Results

### Primary cilia defects in striatal cholinergic interneurons of G2019S LRRK2 mice

We showed previously that cholinergic interneurons that represent about 5% of the neurons in the dorsal striatum of 7-month, R1441C LRRK2 knock-in (KI) mice have fewer primary cilia than their wild type littermates (Dhekne et al., 2018). G2019S LRRK2 is the most common PD-associated mutation in humans, thus it was also important to investigate ciliation in the brains of G2019S LRRK2 mice. G2019S LRRK2 is a hyperactive kinase but less active than R1441C LRRK2 when assayed in cell culture (cf. Steger et al., 2016). The age of the mice was also important: mouse cilia lengthen with age (Arellano et al., 2012) and LRRK2 mutations are incompletely penetrant, making age an important variable in disease onset (Lee et al., 2017; Domingo et al., 2018).

Two different mouse lines were examined: 13-month C57BL/6J mice carrying a genetic knock-in of human LRRK2 G2019S (hereafter referred to as G2019S LRRK2 KI; Steger et al., 2016) and 10-month C57BL/6J mice overexpressing a BAC transgene encoding human G2019S LRRK2 (hereafter referred to as G2019S LRRK2 BAC Tg; Li et al., 2010). Similar to what we observed previously for 7-month R1441C LRRK2 mice, 13-month G2019S LRRK2 KI mice also displayed a significant loss of primary cilia in choline acetyltransferase (ChAT) interneurons of the dorsal striatum (**Figure 1A,B**). As before, primary cilia loss was cell-type specific as there was no difference in ciliation or cilia length in the surrounding cells (primarily medium spiny neurons) between mutant and wild type groups (**Figure 1 – figure supplement 1**). For cholinergic neurons that retained cilia, we detected no significant difference in length between wild type and mutant groups (**Figure 1C**). Cilia length is important to assess as it is thought to reflect signaling capacity (Guo et al., 2017).

**Fig 1.**
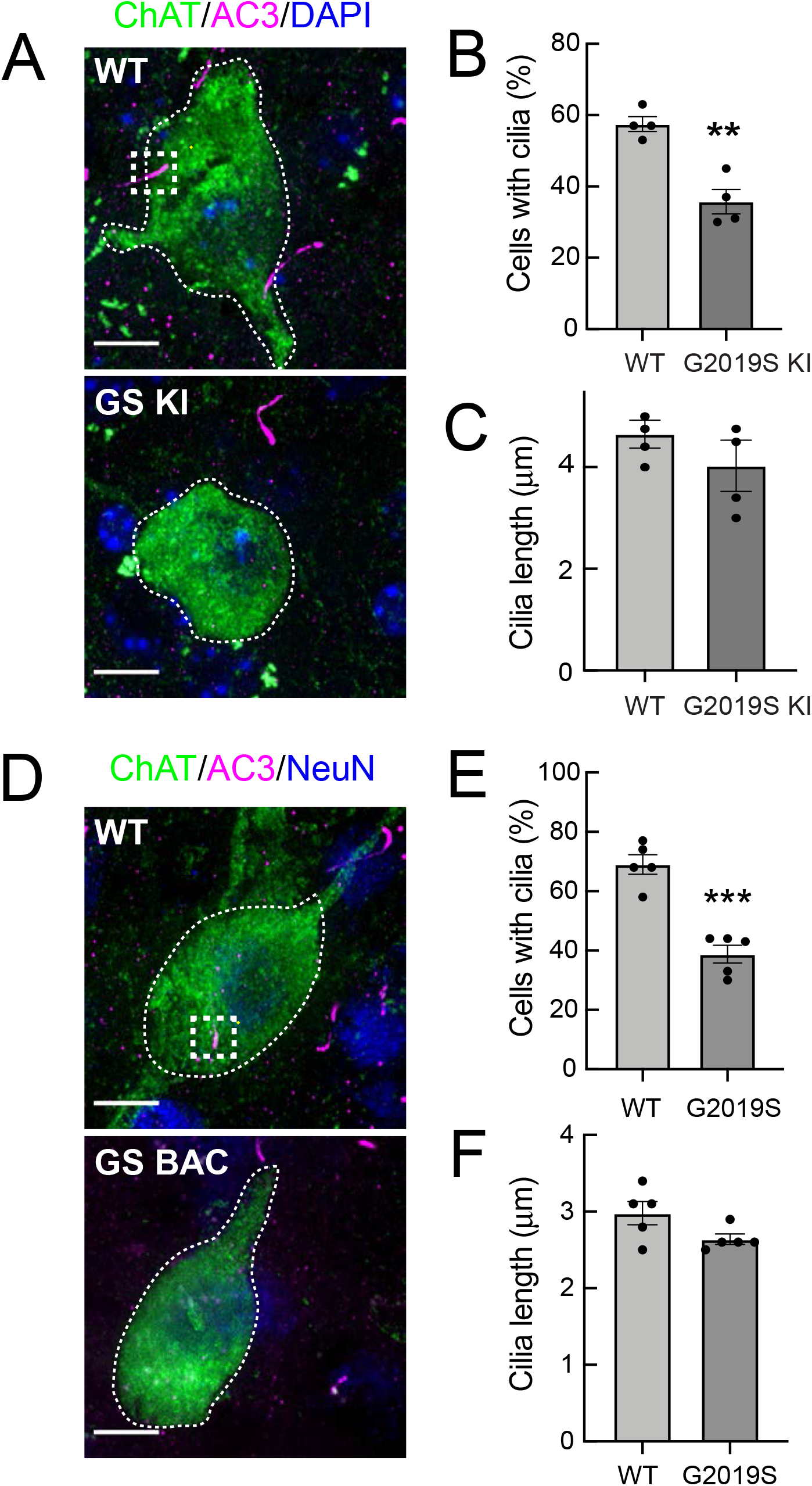
G2019S LRRK2 Striatal Cholinergic Interneurons have fewer primary cilia. **A**. Confocal images of sections of the dorsal striatum from 13-month G2019S LRRK2 KI mice; Cholinergic Acetyltransferase (ChAT) (green, white dotted outline); Adenylate cyclase 3 (AC3) (magenta, white dashed box), and DAPI (blue). **B**. Percentage of ChAT^+^ neurons containing a cilium. Wild type, light gray; G2019S KI, dark gray as indicated. **C**. Quantitation of ChAT^+^ neuron ciliary length from sections as in A. **D**. Confocal images of sections of the dorsal striatum of 10-month G2019S LRRK2 BAC Tg mice; ChAT (green, white dotted outline), AC3 (magenta, white dashed box), and neuron specific nuclear antigen, NeuN (blue). **E**. Percentage of G2019S LRRK2 BAC or wild type ChAT^+^ interneurons containing a cilium. **F**. Quantitation of ChAT^+^ neuron ciliary length. Scale bar, 10 µm. Significance was determined by t-test; (B) **, P = 0.0016; (E) ***, P = 0.0001. Values represent the data from individual brains, analyzing 4-5 brains per group, 2-3 sections per mouse, and >30 neurons per mouse.

G2019S LRRK2 BAC Tg mice overexpress LRRK2 protein by approximately six-fold (Li et al., 2010) and might be expected to display a more severe phenotype than the G2019S LRRK2 KI mice. In the brains of the 10-month G2019S LRRK2 BAC Tg mice we also detected a decrease in ciliated cholinergic interneurons (**Figure 1D**,**E**), comparable to that seen in the G2019S KI mice; when a primary cilium was present, it was not significantly shorter than wild type control cells (**Figure 1F**); again, there was no change in ciliation for the surrounding neurons. Taken together, these data show that G2019S LRRK2 mice have primary cilia defects of the same magnitude as previously observed in cholinergic neurons of the dorsal striatum of R1441C LRRK2 KI mice and cultured cells (Steger et al., 2017; Dhekne et al., 2018). Unlike the R1441C mice however, G2019S mice did not show defects in cortical ciliogenesis.

### Ciliary defects in G2019S Striatal Astrocytes

Mouse LRRK2 is more highly expressed in astrocytes than neurons (cf. Zhang et al., 2014; http://www.brainrnaseq.org), thus it was important to explore the consequences of LRRK2 mutation on mouse astrocyte ciliation. We used glial fibrillary acidic protein (GFAP) and S100 Calcium Binding Protein B (S100B) to identify astrocytes in the striatum. Astrocytic primary cilia were detected using antibodies specific for ADP Ribosylation Factor Like GTPase 13B (Arl13B) because astrocytes in adult mouse brain do not express detectable amounts of Adenylate Cyclase 3 that we use to stain cilia in neurons (Dhekne et al. 2018; Sterpka and Chen, 2018; Sipos et al., 2018; Kasahara et al., 2014). Strikingly, we found that GFAP^+^ G2019S LRRK2 KI and G2019S BAC Tg astrocytes in the dorsal striatum were less likely to have Arl13B^+^ primary cilia relative to wild type controls (**Figure 2**). In addition, the remaining GFAP^+^ cilia in the G2019S BAC Tg astrocytes were very slightly but significantly shorter (**Figure 2F**). Thus, GFAP^+^ striatal astrocytes from G2019S LRRK2 KI mice have fewer primary cilia, and striatal astrocytes also have shorter primary cilia upon pathogenic LRRK2 overexpression. Note that astrocyte cilia were shorter overall compared with neuronal cilia (**Figures 1C**,**F and 2C**,**F**), and all cilia were shorter in the BAC Tg mice.

**Fig 2.**
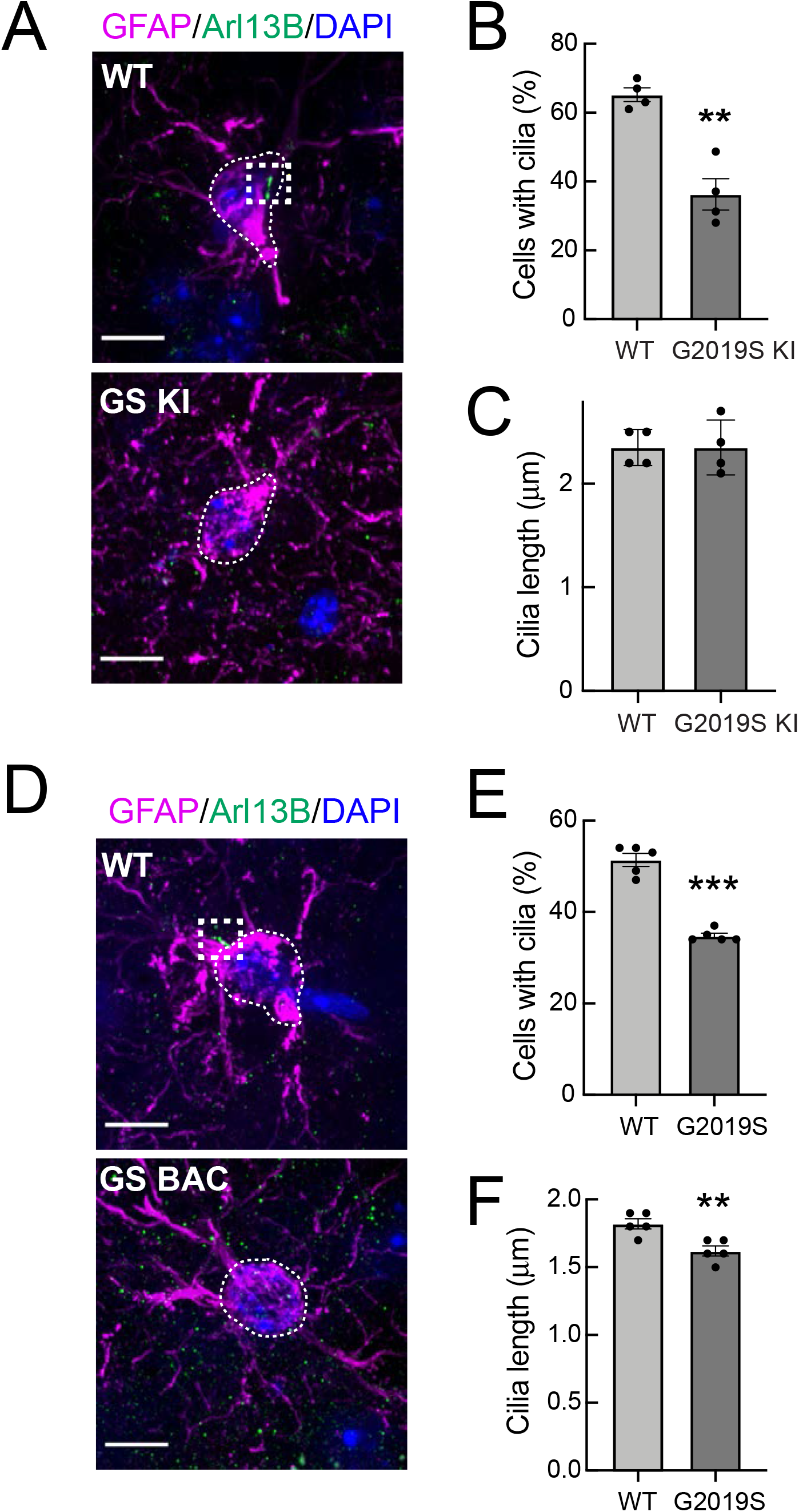
G2019S LRRK2 Striatal Astrocytes have fewer primary cilia. **A**. Confocal images of sections of the dorsal striatum from 13-month G2019S LRRK2 KI mice; astrocyte marker, Glial fibrillary acidic protein (GFAP) (magenta, white dotted outline), cilia marker, ADP-ribosylation factor-like protein 13B (Arl13B) (green, white dashed box), and DAPI (blue). **B**., **C**., Quantitation of the percentage of astrocytes containing a cilium and astrocyte ciliary length from sections described in A. **D**. Confocal images of sections of the dorsal striatum from 10-month G2019S LRRK2 BAC Tg mice, antibody labeled for Glial fibrillary acidic protein (GFAP, magenta, white dashed box), Arl13B (green, white box), and stained with DAPI (blue). **E, F**. Quantitation of the percentage of astrocytes containing a cilium and astrocyte ciliary length from sections described in D. Scale bars, 10 µm. Values represent 4-5 brains per group, 2-3 sections per mouse, and >30 astrocytes per mouse. Significance was determined by t-test; B, **, P = 0.0011; E, ** P = 0.0054; F, **** P < 0.0001.

As shown in **Figure 3A**,**B**, primary cilia loss in cholinergic interneurons of the dorsal striatum could be detected as early as 10 weeks of age in R1441C LRRK2 KI mice. At this age, fewer striatal cholinergic interneurons have a primary cilium (∼40%) in comparison with their non-transgenic littermates (∼60%; **Figure 3B**). Overall neuronal ciliation in the dorsal striatum in 10-week wild type and R1441C mutant groups was slightly lower but similar to the overall values seen previously in a 7-month R1441C LRRK2 KI cohort (∼70%; Dhekne et al 2018).

**Fig 3.**
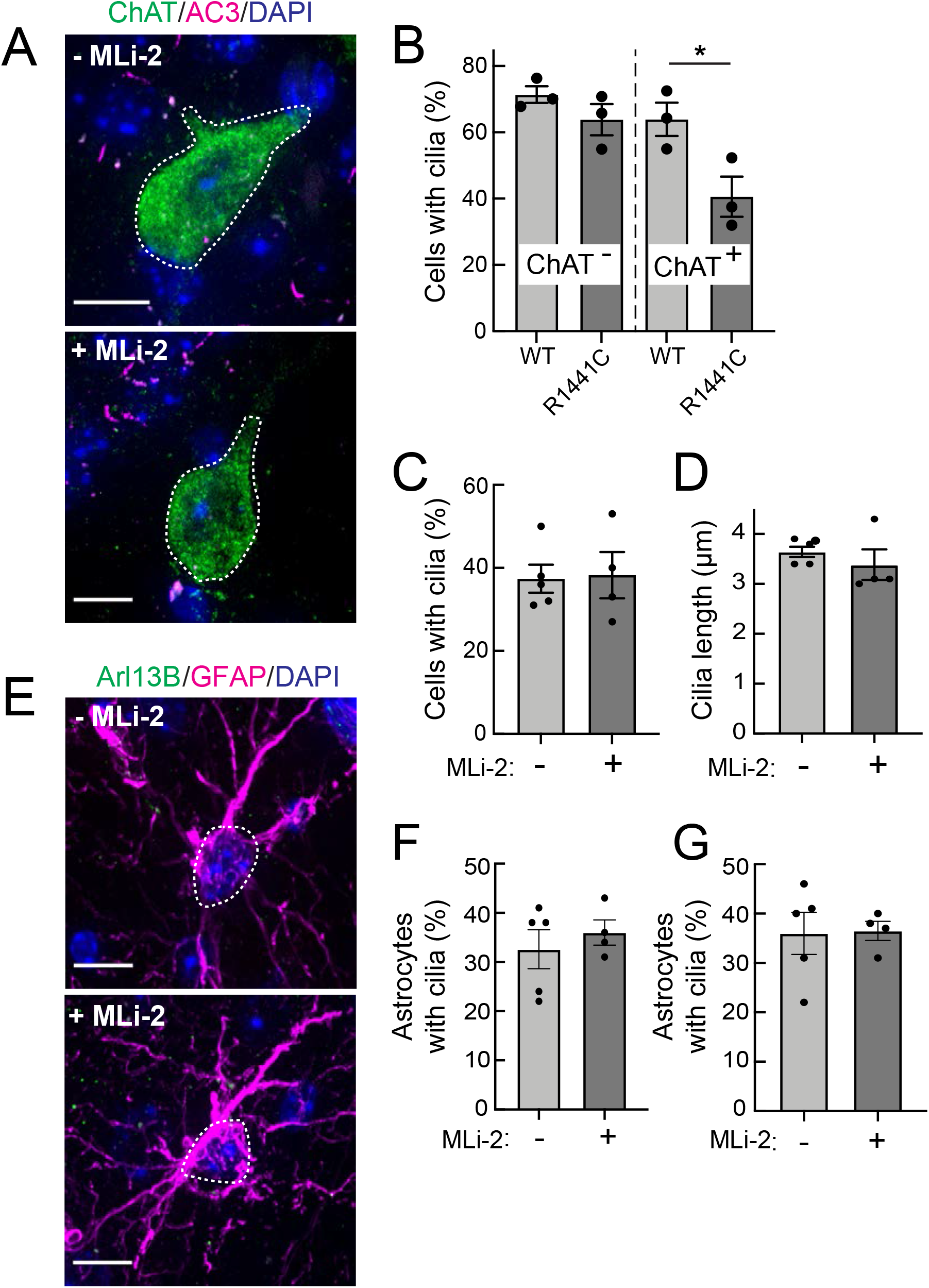
Two weeks MLi-2 treatment does not alter ciliogenesis in R1441C LRRK2 striatal interneurons or astrocytes. Mice (8-weeks old) were fed MLi-2 LRRK2 inhibitor-containing chow or control chow for two consecutive weeks prior to perfusion and staining. **A**. Confocal images of sections of the dorsal striatum from 8 week R1441C LRRK2 KI mice; ChAT (green, white dotted outline); Adenylate cyclase 3 (AC3) (magenta), and DAPI (blue). **B**. Quantitation of the percentage of ChAT^+^ and ChAT^-^ neurons containing a cilium. **C**. Percentage of ChAT^+^ neurons containing a cilium ± MLI-2. **D**. Quantitation of ChAT^+^ neuron ciliary length. **E**. Confocal images of Astrocytes identified by antibodies to GFAP (magenta, white dotted outline), Arl13B (green), and DAPI (blue), ± MLi-2. **F**., **G**. Quantitation of the percentage of total astrocytes (F) or GFAP/S100B^+^ astrocytes (G) containing a cilium. Scale bars, 10 µm. Significance was determined by t-test. B, *, P =0.0417.

### MLi-2 treatment failed to reverse cilia loss in young R1441C LRRK2 KI mice

LRRK2 kinase inhibitors are of great interest due to their potential to normalize LRRK2 kinase activity in patients carrying hyperactive LRRK2 mutant forms. Moreover, if ciliation is a relevant disease phenotype, it would be important to know if cilia defects can be corrected in mutant mice treated with a LRRK2 inhibitor. R1441C LRRK2 KI mice were fed the LRRK2 inhibitor, MLi-2, for two-weeks for phenotypic analysis.

As shown in **Figure 3A**, **C**,**D**, two weeks of MLi-2 feeding did not alter the extent of ciliation or cilia length in cholinergic interneurons or astrocytes in the LRRK2 R1441C mice. We also scored primary cilia in two distinct populations of astrocytes – cells that were positive for both GFAP and S100B, and those that were only positive for S100B (**Figure 3E-G**). Despite two weeks of MLi-2 administration, we failed to detect a significant difference in ciliation or cilia length for either population of astrocytes upon treatment. It is important to note that neurons and astrocytes are postmitotic cells with much less dynamic cilia than dividing cells (Sterpka and Chen, 2018); longer times of drug treatment may very well be needed to reveal an effect.

The ability of MLi-2 to block LRRK2-mediated Rab10 phosphorylation in whole, R1441C LRRK2 mutant mouse brains was assessed by western blot (**Figure 4A**,**B**). LRRK2 inhibitors decrease LRRK2 phosphorylation at serine 935 (cf. Fell et al., 2015), and pS935-LRRK2 was diminished in wild type and R1441C LRRK2 brains from mice fed MLi-2 inhibitor, consistent with inhibitor access in this tissue. Rab12 S105 phosphorylation also decreased in wild type (**Figure 4C**) and mutant (**Figure 4A**,**B**) brain (**Figure 4**). However, levels of pRab10 seemed unchanged (**Figure 4A**,**B**) with variability among animals (see also Ianotta et al., 2020). Previous studies have also noted that MLi-2 administration does not markedly reduce pRab10 levels in brain (cf. Kalogeropulou et al., 2020; Nirujogi et al., 2021). The persistence of pRab10 might explain the persistent cilia loss phenotype in drug treated mice (**Figure 3**).

**Fig 4.**
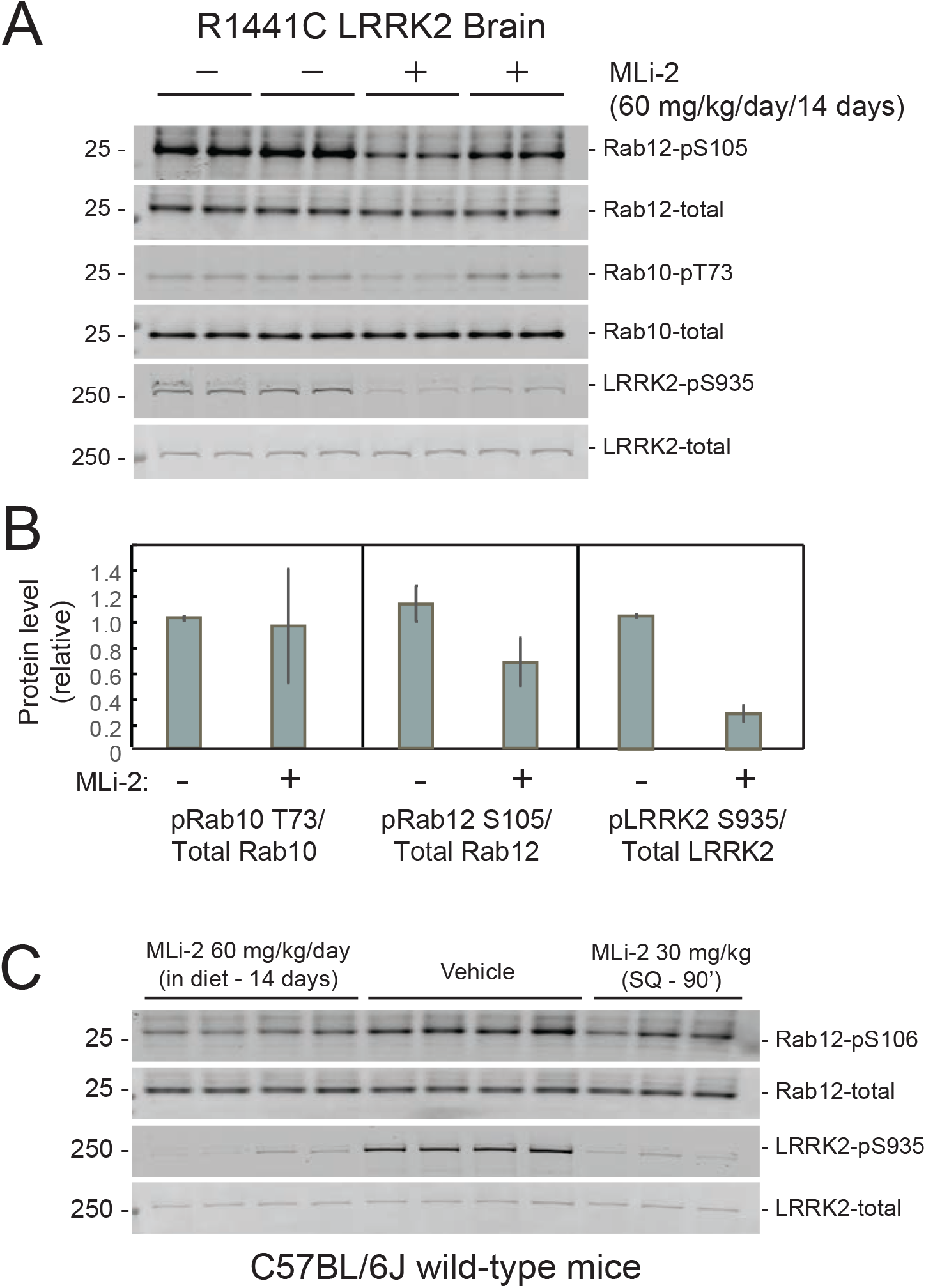
Two weeks MLi-2 treatment decreases LRRK2 pS935 but does not alter Rab10 phosphorylation in R1441C LRRK2 mouse brain. **A** and **B**. Littermate- or age-matched LRRK2 R1441C homozygous knock-in mice were fed either a control diet or MLi-2-containing diet for 14 days prior to tissue collection. Two mice from each group were perfused with PBS; brains were snap frozen in liquid nitrogen and used to monitor inhibition of LRRK2 activity by immunoblotting. **A**. 40 µg brain tissue extract was subjected to immunoblot analysis with antibodies specific for the indicated antigens. Duplicate samples were analyzed using the LI-COR Odyssey CLx imaging system. **B**. Quantitation of data in A, calculated using Image Studio software (mean ± SD, normalized to control diet fed animals.) **C**. Eleven C57BL/6j wild-type mice received either control diet or diet containing MLi-2, targeted to provide a concentration of 60 mg/kg per day for 14 days. On the last day, 3 mice from the control diet group received 30mg/kg MLi-2 dissolved in 40% (w/v) (2-hydroxypropyl)-β-cyclodextrin via subcutaneous injection for 2 hrs prior to tissue collection. 40 µg brain tissue extract was subjected to quantitative immunoblotting analysis with the indicated antibodies. Each lane represents a tissue sample from a different animal.

### PPM1H deficiency phenocopies LRRK2 mutation

PPM1H phosphatase specifically dephosphorylates LRRK2 substrates (Berndsen et al., 2019). If LRRK2 action is responsible for loss of cilia in mutant mouse brains, loss of the corresponding phosphatase should yield the same phenotype in that tissue. **Figure 5** shows that even heterozygous loss of PPM1H leads to decreased cilia numbers in cholinergic interneurons (**Figure 5A**,**B**) and astrocytes (**Figure 5C**,**D**) of the dorsal striatum. These data demonstrate the importance of high PPM1H levels in wild type brain to counteract LRRK2 action. More importantly, they strongly validate a link between LRRK2-Rab phosphorylation and ciliogenesis in specific brain cell types. Immunoblotting showed ∼50% reduction of PPMH levels in brain extracts from PPM1H-/+ mice with levels of pRab10 barely changed at least for whole brain tissue (Fig. 5E).

**Fig 5.**
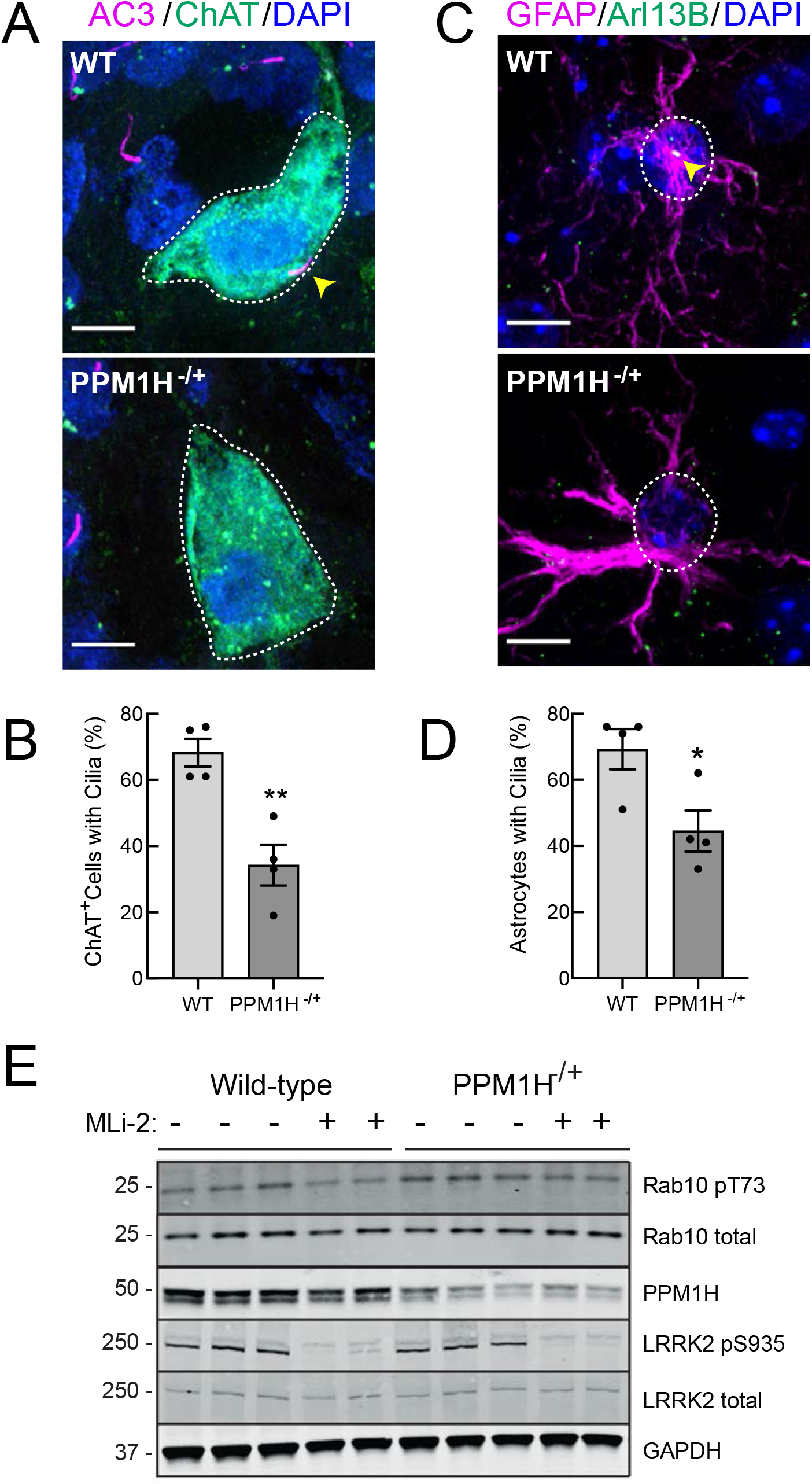
PPM1H^-/+^ Striatal cholinergic neurons and astrocytes have fewer primary cilia. **A**. Confocal images of sections from 6-month PPM1H^-/+^ or age-matched WT mice; ChAT (green, white dotted outline); AC3 (magenta, yellow arrowhead), and DAPI (blue). **B**. Percentage of ChAT^+^ interneurons containing a cilium. Wild type, light gray; PPM1H^-/+^, dark gray as indicated. **C**. Confocal images of sections of the dorsal striatum from 6-month PPM1H^-/+^ or age-matched WT mice; GFAP (magenta, white dotted outline), Arl13B (green, yellow arrowhead), and DAPI (blue). **D**. Percentage of GFAP^+^ astrocytes containing a cilium. Wild type, light gray; PPM1H^-/+^, dark gray as indicated. Scale bars, 10 µm. Significance was determined by t-test; (B) **, P = 0.0058; (D) ***, P = 0.0002. Values represent the data from individual brains, analyzing 4 brains per group, 2-3 sections per mouse, and >30 cells per mouse. **E**. Wildtype and PPM1H^-/+^ knock-out mice were treated with vehicle (40% (w/v) (2-hydroxypropyl)-β-cyclodextrin) or 30 mg/kg MLi-2 dissolved in vehicle by subcutaneous injection 2 hr prior to tissue collection. 40 µg brain tissue extract was analyzed by quantitative immunoblotting analysis with the indicated antibodies. Each lane represents extract from a different mouse.

### Sonic hedgehog signaling is altered in mutant LRRK2 mice

Although Hh signaling is critical during neuronal development, little is known about Hh signaling in the adult brain. As mentioned earlier, striatal ChAT^+^ interneurons respond to Sonic hedgehog ligands that originate from dopaminergic neurons in the substantia nigra (cf. Gonzalez-Reyes et al., 2012). Sonic hedgehog signaling requires cilia: the transmembrane transducer Smoothened must translocate to the primary cilium to initiate the signaling cascade that results in the expression of the transcription factor *Gli1*, a direct Hh target gene that serves as a widely-used metric for signaling strength (Corbit et al, 2005; Rohatgi et al, 2007). Because primary ciliogenesis is likely critical for proper Hh sensing and subsequent GDNF production by cholinergic interneurons, we examined Hh signaling in LRRK2 mutant animals with pathogenic LRRK2 mutations and associated ciliary defects.

*Gli1* transcripts were detected in brain slices using the RNAscope method of fluorescence *in situ* hybridization. **Figure 6A** shows detection of *Gli1* transcripts in wild type, ChAT^+^ neurons, which appear as white dots; a negative control hybridization probe yielded no signal under parallel conditions (**Figure 6A**, bottom row). As expected, ciliated ChAT^+^ neurons showed higher levels of *Gli1* transcripts compared with non-ciliated cells in both mutant and wild type mice (**Figure 6D**). Ciliated cholinergic neurons also displayed the highest number of *Gli1* positive dots compared with all other cell types in the striatum (monitored by DAPI staining) and compared with non-ciliated cholinergic neurons (**Figure 6C**,**D**). This matches well with prior estimates that expression of the Hh receptor PTCH1 is restricted to 6% of total striatal neurons representing all cholinergic and fast spiking interneurons (Gonzalez-Reyes et al., 2012). Importantly, the number of *Gli1* dots was significantly higher in ciliated, striatal cholinergic interneurons of R1441C LRRK2 mice (3-4 *Gli1* positive dots) compared with wild type mice (**Figure 6D**,**E**).

**Fig 6.**
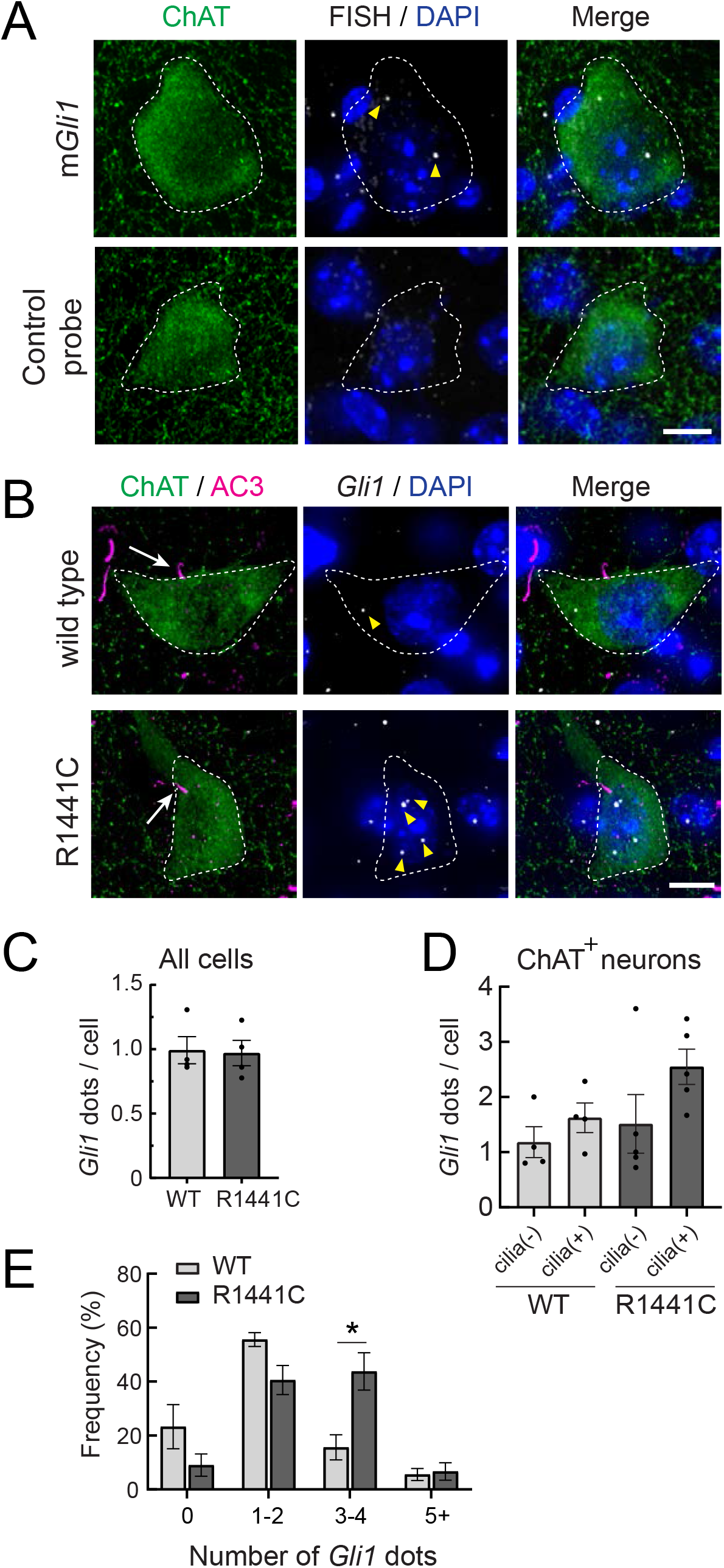
*Gli1* expression in R1441C LRRK2 Dorsal Striatal Cholinergic Interneurons is cilia dependent and enhanced. **A**. 10-month WT mouse dorsal striatum was subjected to *in situ* hybridization using a *Gli1* probe (gray dots, highlighted by yellow arrowheads) or a negative control probe. ChAT (green, white dashed outline) and DAPI^+^ nuclei (blue) were detected by immuno- or chemical staining. **B**. 10-month WT or R1441C mouse dorsal striatum was labeled as indicated: ChAT (green, white dashed outline), AC3 (magenta, white arrow), *Gli1* mRNA (gray dots, yellow arrowheads), DAPI (blue). **C**. Average numbers of *Gli1* dots for all cell types in the dorsal striatum. Cell numbers were determined by DAPI staining. Values represent the mean ± SEM from 4 WT and 4 R1441C brains each containing >500 DAPI stained nuclei from 30 regions. P = 0.88. **D**. Average numbers of *Gli1* dots for cholinergic interneurons with or without primary cilia as indicated. Values represent the mean ± SEM from 4 WT and 5 R1441C brains, each containing 9-32 cells. **E**. Histogram of the number of *Gli1* dots in ciliated cholinergic interneurons from WT or R1441C mice. P = 0.14 (0), 0.054 (1-2),*,0.015 (3-4), 0.79 (5-). Significance was determined by t-test. Arrows indicate primary cilia for ChAT interneurons. Arrowheads indicate *Gli1* mRNA dots. Scale bars, 10µm.

Taken together, these data show that although overall ciliation is decreased, the remaining, ciliated, striatal cholinergic interneurons in R1441C LRRK2 mice transcribe higher quantities of *Gli1* in response to R1441C LRRK2 expression. The upregulation of *Gli1* transcripts in these striatal neurons could be due to increased Hh production in the substantia nigra; perhaps the overall decrease in ciliation decreases GDNF production in the striatum, increasing Hh production by subsequently GDNF-deprived dopaminergic neurons. Alternatively, the remaining cilia in LRRK2 mutant brains may be structurally intact but functionally altered, leading to increased *Gli1* transcript levels. Finally, these data also reveal higher levels of Hh signaling in cholinergic neurons in the dorsal striatum compared with their surrounding neighbors, as predicted by the distribution of PTCH1 protein (Gonzalez-Reyes et al., 2012).

Because R1441C LRRK2 striatal astrocytes also have fewer cilia, we determined if their ability to sense or respond to Hh was also altered. Astrocytes express higher levels of PTCH1 receptor than neurons in both rodent and human brain, and thus would be expected to be capable of Hh signaling. **Figure 7A**,**B** shows astrocytic *Gli1* detection using RNAscope; two classes of astrocytes were scored: those expressing S100B or GFAP. Consistent with the requirement for cilia for canonical Hh signaling, ciliated astrocytes had more *Gli1* dots than non-ciliated astrocytes, independent of whether or not the cells expressed pathogenic R1441C LRRK2 (**Figure 7C**,**D**): in contrast to ChAT^+^ interneurons (**Figure 6**), ciliated and non-ciliated striatal astrocytes from R1441C LRRK2 had similar numbers of *Gli1* positive dots relative to their ciliated wild type controls. Altogether, these data show that striatal astrocytes respond to Hh signaling similarly in R1441C LRRK2 and wild type mice on an individual cell basis, but overall loss of ciliation due to the LRRK2 mutation leads to an overall decrease in striatal astrocytic Hh response.

**Fig 7.**
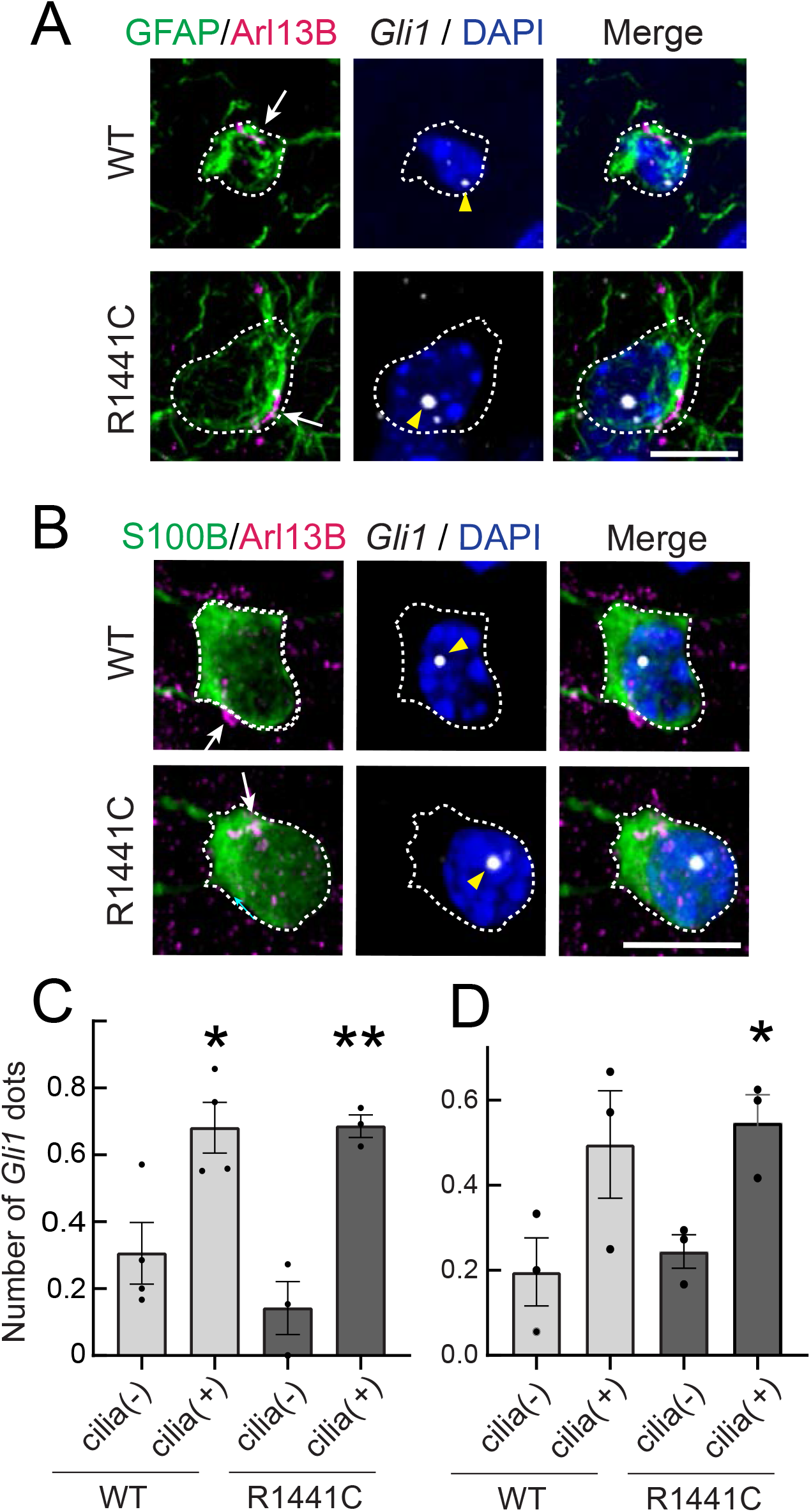
Gli1 expression in astrocytes is cilia dependent. **A**,**B**. 10-month R1441C LRRK2 KI mouse dorsal striatum was subjected to *in situ* hybridization using a *Gli1* probe (gray dots, highlighted with yellow arrowheads). Astrocytes were detected with (A) anti-GFAP or (B) anti-S100B (green, white dashed outline); primary cilia were detected with anti-Arl13B (magenta, white arrows). **C**,**D**. Average numbers of *Gli1* dots from (C) GFAP and (D) S100B^+^ astrocytes with (+) or without (-) primary cilia. Values represent the mean ± SEM from (C) 4 WT and 3 R1441C brains each containing >35 cells (D) 3 WT and 3 R1441C brains each containing >21 cells. (C) WT cilia (-) vs WT cilia (+); *,P =0.020, R1441C cilia (-) vs R1441C cilia (+); **, P =0.0032. (D) WT cilia(-) vs WT cilia(+); P =0.12, R1441C cilia(-) vs R1441C cilia(+); *,P = 0.017. Significance was determined by unpaired t-test. Arrows indicate primary cilia from GFAP or S100B^+^ astrocytes. Scale bars, 10µm.

### Phosphorylated Rab10 in astrocyte ciliogenesis

Our previous work in mouse embryonic fibroblasts and patient-derived iPS cells showed the importance of LRRK2-phosphorylated Rab10 and its effector, RILPL1 in primary ciliogenesis blockade (Dhekne et al., 2018). Unfortunately it was not possible to detect pRab10 in mouse brain sections. To explore further the possible contribution of pRab10 to ciliogenesis blockade in astrocytes, we used immunopanning methods (Foo et al., 2011) to obtain primary, poorly dividing astrocytes from G2019S^+/-^ LRRK2 rat brains. Cells are grown in defined, serum-free media containing 5ng/ml soluble heparin binding EGF-like growth factor (HbEGF) that activates the EGF receptor (Citri and Yarden, 2006) and acts via the EGF receptor in these cells (Foo et al., 2011). Cell density is also relevant as cultured astrocytes secrete other autocrine trophic factors, and ciliogenesis is enhanced in cell culture by increased cell density.

Panned astrocytes showed ∼ 60% ciliation when grown under sparse conditions (**Figure 8A**,**B**) as monitored using anti-Arl13B antibodies. We noted that overnight withdrawal of HbEGF from the growth medium led to ∼50% cilia loss for G2019S^+/-^ LRRK2 astrocytes but not wild type astrocytes under these conditions (**Figure 8B**). Loss of cilia was not seen if HbEGF was withdrawn from wild type cultures or mutant cell cultures in the presence of MLi-2 kinase inhibitor, suggesting that enhanced LRRK2 activity was needed for cilia loss under these culture conditions (**Figure 8B**). We have reported roles for LRRK2 in both cilia formation and cilia loss in other cell types (Sobu et al., 2021). Similar to what we have detected in multiple cell types in culture (Steger et al., 2017; Dhekne et al., 2018), pRab10 levels were highest in non-ciliated cells in the absence of the MLi-2 LRRK2 inhibitor (**Figure 8C**), consistent with a correlation between pRab10 and ciliogenesis.

**Fig 8.**
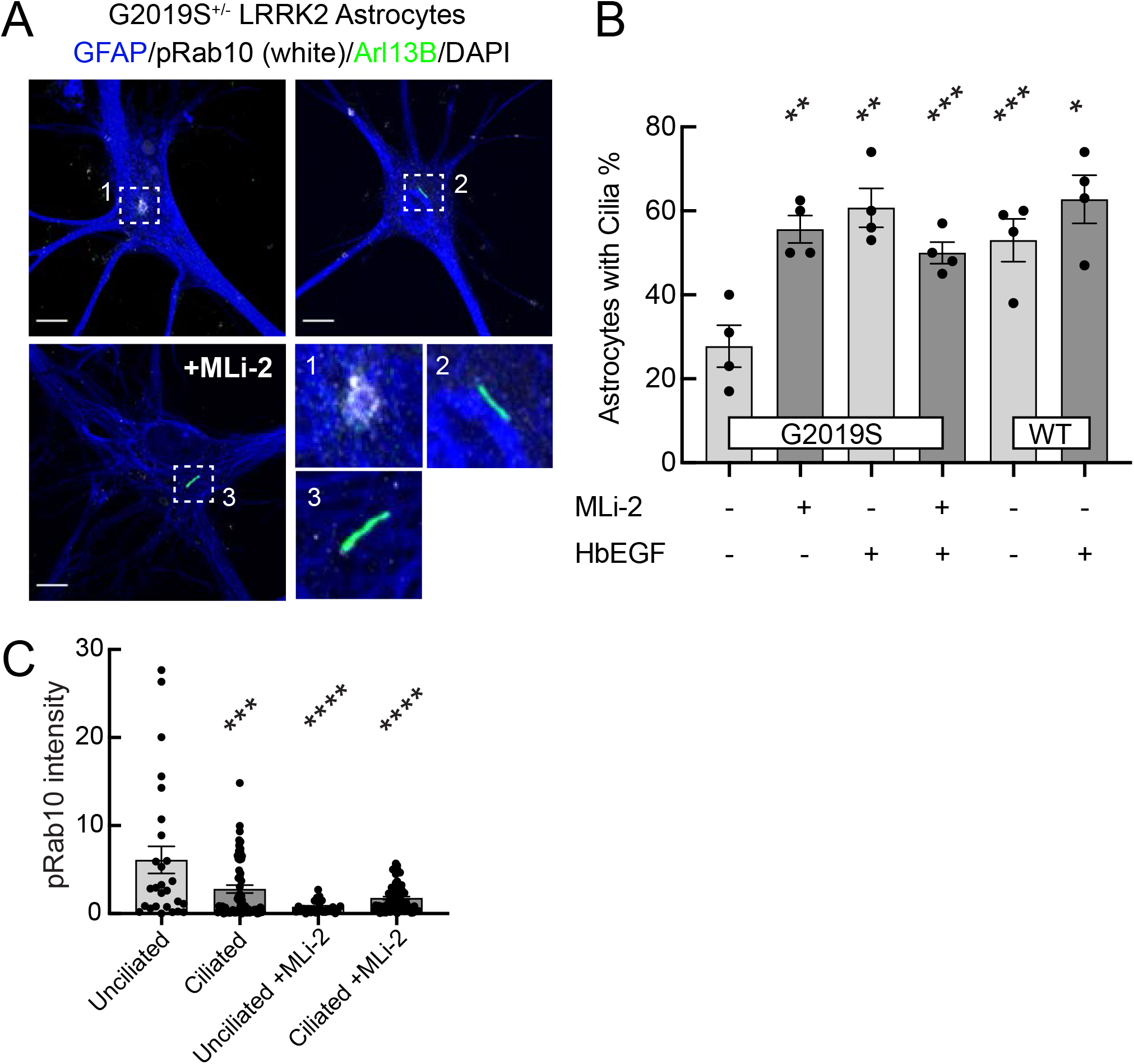
Immuno-panned primary G2019S LRRK2 Astrocytes Display Increased pRab10 and Cilia Loss Upon HbEGF Starvation or AKT Inhibition. BAC Transgenic G2019S^+/-^ LRRK2 rat astrocytes were dissected from P5 pups and cultured for 1 week ± 200 nM MLi-2. **A**. Astrocytes were labeled with anti-GFAP and DAPI (blue); rabbit-anti-phospho-Rab10 (white), and anti-Arl13B (green). The dashed line-boxed, numbered regions were enlarged and are shown at the lower right as numbered. Quantitation of the percent of cells displaying primary cilia in cultures of G2019S^+/-^ or WT astrocytes after 24 hrs ± HbEGF ± 200 nM MLi-2 as indicated. **C**. Quantitation of pRab10 intensity in ciliated or non-ciliated G2019S^+/-^ astrocytes as in B. (B) HbEGF starved G2019S^+/-^ vs. HbEGF starved G2019S^+/-^ plus MLi-2; **, P=0.0017, HbEGF starved G2019S^+/-^ vs. HbEGF fed G2019S^+/-^; **, P=0.0041, HbEGF starved G2019S^+/-^ vs. HbEGF fed G2019S^+/-^ plus MLi-2; ***, P=0.0002, HbEGF starved G2019S^+/-^ vs. HbEGF starved WT; ***, P=0.0003, HbEGF starved G2019S^+/-^ vs. HbEGF fed WT, *, P=0.0112. (C) Non-ciliated G2019S^+/-^ vs Ciliated G2019S^+/-^; ***, P=0.0009, Non-ciliated G2019S^+/-^ vs MLi-2 treated Non-ciliated G2019S^+/-^; ****, P<0.0001; Non-ciliated G2019S^+/-^ vs MLi-2 treated ciliated; ****, P<0.0001; ns = not statistically significant. Significance was determined either by student’s t-test or by Ordinary one-way ANOVA using Dunnett’s multiple comparisons test. Scale bars, 10µm.

## Discussion

Pathogenic LRRK2 activity causes primary cilia loss in striatal cholinergic interneurons in mature R1441C LRRK2 mice (Dhekne et al., 2018). In this study, we show that primary cilia loss is also seen in two distinct G2019S LRRK2 mouse models of the most common human LRRK2 mutation. Moreover, our study reveals that primary cilia loss in cholinergic interneurons can be detected as early as 10 weeks of age. Thus, pathogenic LRRK2 activity causes interneuron primary cilia loss at a relatively early stage and loss is sustained throughout the animal’s lifespan for post-mitotic neurons. Consistent with this conclusion was our finding that mice deficient in the phospho-Rab-specific phosphatase, PPM1H, show the same ciliary defect as mice harboring a pathogenic LRRK2 mutation. This provides strong genetic evidence for the importance of LRRK2 activity in regulating cilia formation, in both LRRK2 wild type/PPM1H deficient and LRRK2 mutant animals.

Striatal cholinergic interneurons provide trophic support to dopaminergic neurons via Hh sensing (Gonzalez-Reyes et al., 2012), and primary cilia loss is predicted to perturb the ability of cholinergic neurons to sense Hh ligands as required for this role. Our FISH results support this model by showing that ciliated striatal ChAT^+^ interneurons express abundant *Gli1* transcripts under normal, physiologic conditions, and represent the cell type showing the greatest level of *Gli1* expression in the dorsal striatum. In the presence of the pathogenic R1441C LRRK2 mutation, we observed significant elevation of *Gli1* transcripts in the smaller number of remaining, ciliated, striatal cholinergic interneurons. One explanation for this increase in *Gli1* mRNA is that dopaminergic neurons in the substantia nigra upregulate Hh production due to stress or lack of GDNF signals from the striatum. Other studies of cortical neurons have shown a role for primary cilia in protection from stresses of ethanol- or ketamine-induced caspase activation and dendritic degeneration (Ishii et al., 2021). Future experiments will seek to monitor changes in Hh production in the substantia nigra that may be associated with LRRK2 mutation.

For the first time we show that LRRK2 G2019S striatal astrocytes are also cilia-deficient and the remaining ciliated cells display a modest increase in *Gli1* transcription relative to non-ciliated astrocytes. These results indicate that fewer striatal astrocytes in LRRK2 G2019S mice are capable of responding to Hh ligands and importantly, the remaining astrocytes that do respond, do not compensate for a more general loss of ciliated astrocytes. How might decreased Hh sensing by astrocytes impact the striatum? Striatal astrocytes functionally interact with distinct networks of medium spiny neurons (Khakh, 2019; Martín et al., 2015), and loss of Hh sensing may perturb these interactions. This is relevant because imbalances in medium spiny neuron activity are known to contribute to the slowness and rigidity of movement that is symptomatic of Parkinson’s disease (Zhai et al., 2018).

Chen et al. (2020) recently reported altered organization of glutaminergic AMPA receptors in cultured striatal neurons from LRRK2 G2019S and R1441C mutant mice. This was accompanied by decreased frequency of miniature excitatory post-synaptic currents in brain slices. It is likely that LRRK2-mediated Rab phosphorylation is in some way responsible for these changes; Rab phosphorylation could of course regulate the trafficking of AMPA receptors. It is also possible that the overall physiological changes are due to loss of cilia from the astrocytes that surround these neurons and support their overall physiology.

Decreased Hh signaling in astrocytes decreases Kir4.1 potassium channel expression in the cerebellum and neocortex and impairs turnover of dendritic spines, accompanied by an increase in neuronal excitability (Farmer et al., 2016; Hill et al., 2019). Kir4.1 expression is markedly decreased in several neurological disorders (cf. Scholl et al., 2009; Inyushin et al., 2010; Gilliam et al., 2014), including in striatal cells of Huntington’s disease mice (Tong et al., 2014; Dvorzhak et al., 2016). Striatal astrocytes may use Kir4.1 channels to modulate medium spiny neuron synaptic transmission by local buffering of potassium ions, as shown for other brain regions (Djukic et al., 2007, Sibille et al., 2014). Thus, primary cilia loss from LRRK2 G2019S striatal astrocytes is likely to dysregulate medium spinal neuron circuitry.

Two weeks of LRRK2 MLi-2 inhibitor administration failed to reverse cilia deficits in LRRK2 mutant animals. This may not be surprising as cilia are likely long lived in post-mitotic neurons and non-reactive astrocytes. Very little is known about the dynamics of cilia in the adult brain, however data from experiments in which a critical ciliary component, IFT88 was inducibly knocked out in adult brain indicated that 3 months was needed to see the consequences of loss of this critical ciliary factor (Bowie and Goetz, 2020). Thus, it is possible that longer feeding regimens will reveal a recovery in striatal ciliation. Another puzzle is why pRab10 levels do not change as significantly as pRab12 and LRRK2 pS935 in the brains of drug treated animals. It is possible that in the brain, pRab10 is greatly stabilized by strong binding to an abundant phosphoRab effector(s). Additional work is needed to resolve this mystery.

Altogether, these data demonstrate a profound effect of LRRK2 hyperactivation on primary ciliogenesis in a population of cholinergic interneurons that play a key role in motor function and in astrocytes that support medium spiny neuron function in the dorsal striatum. Loss of neuroprotection from cholinergic interneurons and astrocytes of the striatum may explain the loss of dopaminergic neurons associated with a disease of aging such as Parkinson’s disease.

## Acknowledgements

This study was funded by the joint efforts of The Michael J. Fox Foundation for Parkinson’s Research (MJFF) [17298 & 6986 (S.R. P. & D.R.A.)] and Aligning Science Across Parkinson’s (ASAP) initiative. MJFF administers the grant (ASAP-000463, S.R.P. & D.R.A.) on behalf of ASAP and itself, and the Medical Research Council [grant no.MC_UU_12016/2 (D.R.A.)] and the pharmaceutical companies supporting the Division of Signal Transduction Therapy Unit (Boehringer-Ingelheim, GlaxoSmithKline, Merck KGaA (D.R.A.). We are especially grateful to Dr. Rajat Rohatgi for his critical input.

## Materials and Methods

### Reagents

MLi-2 LRRK2 inhibitor was synthesized by Natalia Shpiro (MRC Reagents and Services, University of Dundee) and was first described to be a selective LRRK2 inhibitor in previous work (Fell et al., 2015).

### Transgenic Mice

All animal studies were performed in accordance with the Animals (Scientific Procedures) Act of 1986 and regulations set by the University of Dundee, the U.K. Home Office, and the Administrative Panel on Laboratory Animal Care at Stanford University. Mice were maintained under specific pathogen-free conditions at the University of Dundee (U.K.); they were multiply housed at an ambient temperature (20–24°C) and humidity (45–55%) and maintained on a 12 hr light/12 h dark cycle, with rodent diet and water available ad libitum. LRRK2 R1441C knock-in mice were obtained from the Jackson laboratory (Stock number: 009346). LRRK2 G2019S knock-in and PPM1H knock-out mice were obtained from Taconic (Model 13940 and TF3142, respectively). G2019S BAC transgenic brain sections from 10-month mice were a gift from Aaron Gitler at Stanford. Mouse genotyping was performed by PCR using genomic DNA isolated from tail clips or ear biopsies.

For the experiment shown in Figure 5E, 63-83 day old mice of the indicated genotypes were injected subcutaneously with vehicle [40% (w/v) (2-hydroxypropyl)-β-cyclodextrin (Sigma–Aldrich #332607)] or MLi-2 dissolved in the vehicle at a 30 mg/kg final dose. Mice were euthanized by cervical dislocation 2hr following treatment and the collected tissues were rapidly snap frozen in liquid nitrogen.

### Mouse brain processing

Homozygous LRRK2-mutant (R1441C or G2019S of various ages as indicated) or heterozygous PPM1H^-/+^ KO mice (164 days old) and age-matched wild-type controls were fixed by transcardial perfusion using 4% paraformaldehyde (PFA) in PBS as described in Khan et al. (2020). Next, whole brain tissue was extracted, post-fixed in 4% PFA for 24 hrs and then immersed in 30% (w/v) sucrose in PBS until the tissue settled to the bottom of the tube (∼48 hrs). LRRK2 R1441C KI, LRRK2 G2019S KI and PPM1H^-/+^ KO brains were harvested in Dundee and sent with identities blinded until analysis was completed. Prior to cryosectioning, brains were embedded in cubed-shaped plastic blocks with OCT (BioTek, USA) and stored at −80°C. OCT blocks were allowed to reach −20°C for ease of sectioning. The brains were oriented to cut sagittal or coronal sections on a cryotome (Leica CM3050S, Germany) at 16-25 µm thickness and positioned onto SuperFrost plus tissue slides (Thermo Fisher, USA).

### In-diet MLi-2 administration to wild-type mice (pilot study) and LRRK2 R1441C KI mice

For the pilot experiment shown in Figure 4C, 11 C57BL/6j wild-type mice were allowed to acclimate to the control rodent diet (Research Diets D01060501; Research Diets, New Brunswick, NJ) for 14 days before being placed on study. On day 1 of the study, one group (4 mice) received modified rodent diet (Research Diets D01060501) containing MLi-2 and formulated by Research Diets to provide a concentration of 60 mg/kg per day on the basis of an average food intake of 5 g/day for 14 days; the other group (7 mice) received untreated diet (Research Diets D01060501) for 14 days and served as the control group. The dose of MLi-2 and the length of the in-diet treatment used for this study were based on Fell et al., 2015. Bodyweight and food intake were assessed twice weekly. On the last day of the study, 3 mice from the control diet group received 30mg/kg MLi-2 dissolved in 40% (w/v) (2-hydroxypropyl)- β-cyclodextrin via subcutaneous injection for 2 hr prior to tissue collection. All mice were euthanized by cervical dislocation and the collected tissues were rapidly snap frozen in liquid nitrogen and brain samples used for quantitative immunoblotting analysis of phospho-Ser935 LRRK2 and phospho-Ser105 Rab12 as a readout of LRRK2 activity.

For the experiments described in Figure 3C-G, littermate or age-matched male and female LRRK2 R1441C homozygous knock-in mice at 10 weeks of age were used. Mice were allowed to acclimate to the control rodent diet for 14 days as described above before being placed on study. On day 1 of the study, one group (7 mice) received a modified rodent diet targeted to provide a concentration of 60 mg/kg per day on the basis of an average food intake of 5 g/day; the other group (7 mice) received an untreated diet and served as the control group. Bodyweight and food intake were assessed twice weekly. On day 15, mice were terminally anesthetized and brains harvested and fixed as described above. For immunoblotting analysis to confirm LRRK2 inhibition, brains were snap frozen in liquid nitrogen from mice that were terminally anesthetized and perfused by injection of PBS into the left cardiac ventricle (2 mice from each group).

### Preparation of mouse tissue lysates for immunoblotting analysis

Snap frozen tissues were weighed and quickly thawed on ice in a 10-fold volume excess of ice-cold lysis buffer containing 50 mM Tris–HCl pH 7.4, 1 mM EGTA, 10 mM 2-glycerophosphate, 50 mM sodium fluoride, 5 mM sodium pyrophosphate, 270 mM sucrose, supplemented with 1 µg/ml microcystin-LR, 1 mM sodium orthovanadate, complete EDTA-free protease inhibitor cocktail (Roche), and 1% (v/v) Triton X-100. Tissues were homogenized using a POLYTRON homogenizer (KINEMATICA), employing three rounds of 10 s homogenization with 10 s intervals on ice. Lysates were clarified by centrifugation at 20 800***g*** for 30 min at 4°C and supernatant was collected for subsequent protein quantification by Bradford assay and immunoblot analysis.

### Quantitative immunoblotting analysis

40 µg of brain extracts were loaded onto a NuPAGE 4–12% Bis–Tris Midi Gel (Thermo Fisher Scientific, Cat# WG1402BOX) and electrophoresed at 130 V for 2 hrs with NuPAGE MOPS SDS running buffer (Thermo Fisher Scientific, Cat# NP0001-02). Proteins were then electrophoretically transferred onto a nitrocellulose membrane (GE Healthcare, Amersham Protran Supported 0.45 µm NC) at 100 V for 90 min on ice in transfer buffer (48 mM Tris–HCl and 39 mM glycine supplemented with 20% (v/v) methanol). The transferred membrane was blocked with 5% (w/v) skim milk powder dissolved in TBS-T (50 mM Tris base, 150 mM sodium chloride (NaCl), 0.1% (v/v) Tween 20) at room temperature (RT) for 30 min and incubated overnight at 4°C in primary antibodies diluted in 5% (w/v) BSA in TBS-T. After incubation with primary antibodies, membranes were washed three times for 5 min with TBS-T and incubated with near-infrared fluorescent dye-labelled secondary antibodies (diluted to 1:20,000) for 1 hr at RT. Thereafter, membranes were extensively washed with TBS-T and protein bands were acquired via near-infrared fluorescent detection using the Odyssey CLx imaging system and the signal intensity quantified using Image Studio software.

### Fluorescence in situ hybridization (FISH)

Wild type (WT) and R1441C LRRK2 KI 10-month mouse brains were sliced coronally at 25 µm thickness. RNAscope® Multiplex Fluorescent Detection Kit v2 (Advanced Cell Diagnostics) was carried out according to the manufacturer using RNAscope® 3-plex Negative Control Probe (#320871) or Mm-*Gli1* (#311001). Opal 690 (Akoya Biosciences) was used for fluorescent visualization of hybridized probes. Then, brain slices were blocked with 0.1% BSA and 10% donkey serum in TBS (Tris buffered saline) containing 0.1% Triton X-100 for 30 mins followed by incubation with primary antibody in TBS + 0.1% BSA and 1% DMSO overnight at 4 °C. Secondary antibody was also diluted in TBS + 0.1% BSA and 1% Triton X-100 for 30 mins and then added for 2 hrs at RT. Sections were mounted with ProLong™ Gold Antifade Mountant with DAPI and glass coverslips.

### Immunofluorescence staining and microscopy

Brain primary cilia detection and analyses were performed as previously described (Khan et al., 2020). Briefly, frozen slides were thawed at RT for 10 min then gently washed with PBS for 5 min. Sections were permeabilized with 0.1% Triton X-100 in PBS at RT for 15 min. Sections were blocked with 2% BSA in PBS for 2 hrs at RT and were then incubated overnight at 4°C with primary antibodies. The following day, sections were incubated with secondary antibodies at RT for 2 hr. Donkey Highly cross absorbed H + L secondary antibodies (Life Technologies) conjugated to Alexa 488, Alexa 568 or Alexa 647 were used at a 1:1000 dilution. Stained tissues were overlayed with Mowiol and a glass coverslip.All antibody dilutions for tissue staining included 1% DMSO to help antibody penetration. All images were obtained using a spinning disk confocal microscope (Yokogawa) with an electron multiplying charge coupled device (EMCCD) camera (Andor, UK) and a 100X 1.4 NA oil immersion objective. All image visualizations and analyses were performed using Fiji (https://fiji.sc). Cholinergic neurons in the dorsal striatal region were identified in the caudate putamen of sagittal or coronal sections that were reactive for both choline acetyltransferase and the pan neuronal marker, NeuN. Dorsal striatal astrocytes were defined as GFAP^+^ and/or S100B^+^ cells in the caudate putamen.

### Astrocyte Immune-Panning and Microscopy

Rat Astrocyte Immune-panning was performed as previously described (Foo et al., 2013; Dhekne et al., 2018). Cells on coverslips were fixed with 3.5% PFA for 15 minutes at RT. The cells were then subjected to three washes with PBS for 5 min each. To permeabilize, samples were incubated with PBS containing 0.1% Saponin for 15 min. All subsequent steps contained 0.05% Saponin unless otherwise specified. Samples were again washed three times with PBS then incubated in blocking solution (PBS containing 2% BSA) for 1 hr at RT. Afterwards, the cells were incubated in blocking solution containing primary antibodies for 1 hr at RT (EnCOR Chicken-anti-GFAP at 1:2000, Abcam Rabbit-anti-pRab10 at 1:1000, Neuromab Mouse-anti-Arl13B at 1:2000). Cells were then washed three times with blocking solution, then incubated for 1 hr at RT with blocking solution containing DAPI and donkey-anti alexa fluor secondary antibodies. Coverslips were then rinsed without saponin in PBS two times, in ddH_2_O once, and then mounted with mowiol. All image visualizations and analyses were performed using Fiji (https://fiji.sc). Maximum intensity projections, background subtraction, and pRab10 integrated intensity measurements were obtained using CellProfiler (https://cellprofiler.org/).

### Statistics

Graphs were made using Graphpad Prism 6 software. Error bars indicate SEM. Unless otherwise specified, a Student’s T-test was used to test significance. Brains were harvested in Dundee and analyzed at Stanford; identities were blinded for unbiased analyses.

## KEY RESOURCES. Primary antibodies and reagents used in this study

| KEY RESOURCES TABLE |  |  |  |  |
| --- | --- | --- | --- | --- |
| Reagent type (species) or resource | Designation | Source or reference | Identifiers | Additional information |
| Genetic reagent<br>(*Rattus norvegicus*) | SA Sprague Dawley rat | Taconic | #SD-M | NTac:SD Background |
| Genetic reagent<br>(*Rattus norvegicus*) | Human LRRK2 G2019S rat | Taconic | #10681 | NTac:SD Background; BAC Transgene |
| Genetic reagent<br>(*Mus musculus*) | Tg(LRRK2*G2019S)2AMjf | Jackson Laboratory | #018785 | C57BL/6 Background; BAC Transgene |
| Genetic reagent<br>(*Mus musculus*) | Constitutive KI<br>*Lrrk2*<sup>tm4.1Arte</sup> | Taconic | #13940 | C57BL/6 Background; G2019S KI |
| Genetic reagent<br>(*Mus musculus*) | B6.Cg-*Lrrk2*<sup>tm1.1Shn</sup>/J | Jackson Laboratory | #009346 | C57BL/6 Background; R1441C KI |
| Genetic reagent<br>(*Mus musculus*) | Constitutive KO mouse | Taconic | #TF3142 | C57BL/6 Background; PPM1H KO |
| Antibody | anti-Arl13B (mouse monoclonal) | Neuromab | N295B/66 | (1:1000) |
| Antibody | anti-Adenylate cyclase III (rabbit polyclonal) | Santa Cruz | SC-588 | (1:100) |
| Antibody | anti-Adenylate cyclase III (rabbit polyclonal) | EnCOR | RPCA-ACIII | (1:10000) |
| Antibody | anti-NeuN<br>(mouse monoclonal) | Proteintech | 66836-1-IG | (1:500) |
| Antibody | anti-Choline<br>Acetyltransferase<br>(goat polyclonal) | Millipore | AB144P-1ML | (1:100) |
| Antibody | anti-GFAP<br>(chicken polyclonal) | Novus<br>Biologicals | NBP1-05198 | (1:2000) |
| Antibody | anti-GFAP<br>(chicken polyclonal) | EnCOR | CPCA-<br>GFAP | (1:2000) |
| Antibody | anti-S100B<br>(guinea pig polyclonal) | Synaptic<br>Systems | #287004 | (1:200) |
| Antibody | anti-phospho-Rab10 (Thr73)<br>(rabbit monoclonal) | Abcam | AB230261 | (1:1000) |
| Antibody | anti-Rab10<br>(mouse monoclonal) | Nanotools | 0680–<br>100/Rab10<br>-605B11 | 1 mg/ml |
| Antibody | anti-phospho-Rab12 (Ser105)<br>(rabbit monoclonal) | Abcam | ab256487 | 1 mg/ml |
| Antibody | anti-Rab12<br>(sheep polyclonal) | MRC PPU<br>Reagents<br>and<br>Services,<br>University of<br>Dundee | SA227 | 1 mg/ml |
| Antibody | anti-phospho-LRRK2 (Ser935)<br>(rabbit monoclonal) | MRC PPU<br>Reagents<br>and<br>Services,<br>University of<br>Dundee | UDD2 | 1 mg/ml |
| Antibody | Anti-LRRK2<br>(mouse monoclonal) | Antibodies<br>Inc./NeuroM<br>ab | 75-253 | 1 mg/ml |
| Antibody | anti-PPM1H<br>(sheep polyclonal) | MRC PPU<br>Reagents<br>and<br>Services,<br>University of<br>Dundee | DA018 | 1 mg/ml |
| Chemical<br>compound,<br>drug | MLi-2 | MRC PPU<br>Reagents<br>and<br>Services,<br>University of<br>Dundee |  |  |
| Commercial assay, kit | RNAscope®<br>Multiplex<br>Fluorescent<br>Reagent Kit v2 | Advanced<br>Cell<br>Diagnostics | #323100 |  |
| Commercial assay, kit | RNAscope® 3-plex<br>Negative Control<br>Probe | Advanced<br>Cell<br>Diagnostics | #320871 |  |
| Commercial assay, kit | RNAscope®<br>Probe<br>- Mm-*Gli1* | Advanced<br>Cell<br>Diagnostics | #311001 |  |
| Commercial assay, kit | OPAL 690<br>REAGENT<br>PACK | Akoya<br>Biosciences | FP1497001K<br>T | (1:750) |
| Rodent diet |  | Research<br>Diets, Inc. | D01060501 | Untreated diet |
| Rodent diet containing MLi-2 |  | Research<br>Diets, Inc. | D01060501<br>added with<br>360 mg MLi-2<br>per kg | Modified diet |
| Software, Algorithm | FIJI | PMID:<br>29187165 |  |  |
| Software, Algorithm | CellProfiler | PMID:<br>29969450 |  |  |

---

**Figure Figure Supplement 1.**
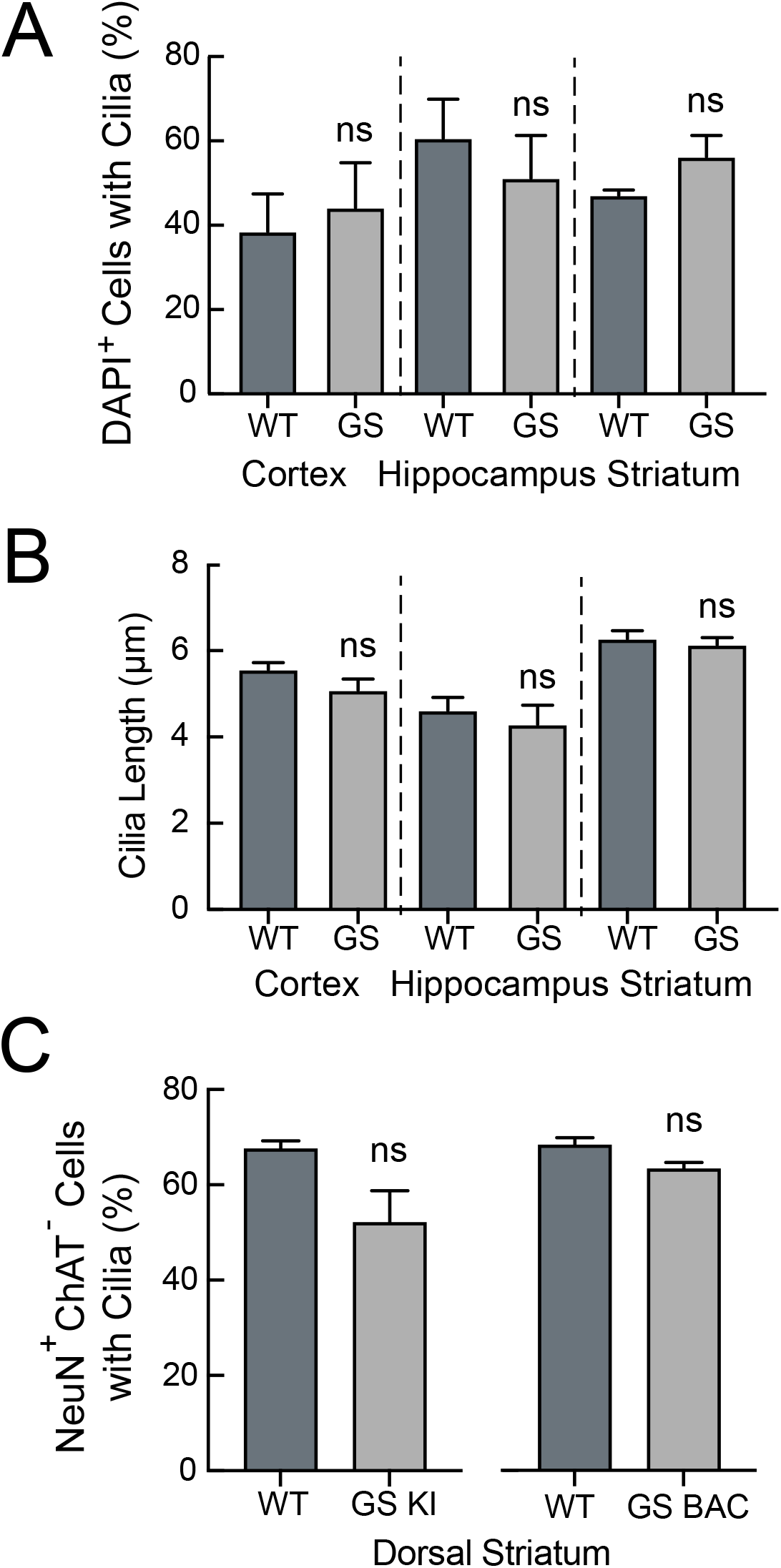
Cilia Density and Cilia Length in G2019S LRRK2 and Wild Type Mice. **A**. Quantitation of cilia density for all cells marked by DAPI staining in the cortex, hippocampus and striatum (as indicated) of 13-month G2019S LRRK2 KI (GS KI) or wild type (WT) mice. **B**. Quantitation of cilia length for DAPI stained cells in the cortex, hippocampus and striatum of 13-month GS KI or WT mice. **C**. Quantitation of NeuN^+^ neurons with cilia in the dorsal striatum of 13-month GS KI and WT mice (left bars) or 10-month GS BAC and WT mice (right bars). Significance was determined by t-test for 2-3 tissue sections per genotype; >50 cells per brain region.

## References

Alessi DR, Sammler E. 2018. LRRK2 kinase in Parkinson’s disease. Science 360(6384):36–37.

Arellano JI, Guadiana SM, Breunig JJ, Rakic P, Sarkisian MR. 2012. Development and distribution of neuronal cilia in mouse neocortex. J Comp Neurol 520:848–873.

Berndsen K, Lis P, Yeshaw WM, Wawro PS, Nirujogi RS, Wightman M, Macartney T, Dorward M, Knebel A, Tonelli F, Pfeffer SR, Alessi DR. 2019. PPM1H phosphatase counteracts LRRK2 signaling by selectively dephosphorylating Rab proteins. Elife 8:e50416.

Bowie E, Goetz SC. 2020. TTBK2 and primary cilia are essential for the connectivity and survival of cerebellar Purkinje neurons. Elife 2020 9:e51166.

Chen C, Soto G, Dumrongprechachan V, Bannon N, Kang S, Kozorovitskiy Y, Parisiadou L. 2020. Pathway-specific dysregulation of striatal excitatory synapses by LRRK2 mutations. Elife 2020 Oct 2;9:e58997.

Citri A, Yarden Y. 2006. EGF-ERBB signalling: towards the systems level. Nat Rev Mol Cell Biol. 7(7):505–16.

Corbit KC, Aanstad P, Singla V, Norman AR, Stainier DYR, Reiter JF. 2005. Vertebrate Smoothened functions at the primary cilium. Nature 437:1018–1021.

Dhekne HS, Yanatori I, Gomez RC, Tonelli F, Diez F, Schüle B, Steger M, Alessi DR, Pfeffer SR. 2018. A pathway for Parkinson’s disease LRRK2 kinase to block primary cilia and sonic hedgehog signaling in the brain. Elife 7:e40202.

Djukic B, Casper KB, Philpot BD, Chin L-S, McCarthy KD. 2007. Conditional Knock-Out of Kir4.1 Leads to Glial Membrane Depolarization, Inhibition of Potassium and Glutamate Uptake, and Enhanced Short-Term Synaptic Potentiation. J Neurosci 27:11354–11365.

Domingo A, Klein C. 2018. Genetics of Parkinson disease, Handbook of Clinical Neurology.

Dvorzhak A, Vagner T, Kirmse K, Grantyn R. 2016. Functional Indicators of Glutamate Transport in Single Striatal Astrocytes and the Influence of Kir4.1 in Normal and Huntington Mice. J Neurosci 36:4959– 4975.

Farmer WT, Abrahamsson T, Chierzi S, Lui C, Zaelzer C, Jones EV, Bally BP, Chen GG, Théroux JF, Peng J, Bourque CW, Charron F, Ernst C, Sjöström PJ, Murai KK. 2016. Neurons diversify astrocytes in the adult brain through sonic hedgehog signaling. Science 351(6275):849–54.

Fell MJ, Mirescu C, Basu K, Cheewatrakoolpong B, DeMong DE, Ellis JM, Hyde LA, Lin Y, Markgraf CG, Mei H, Miller M, Poulet FM, Scott JD, Smith MD, Yin Z, Zhou X, Parker EM, Kennedy ME, Morrow JA. 2015. MLi-2, a Potent, Selective, and Centrally Active Compound for Exploring the Therapeutic Potential and Safety of LRRK2 Kinase Inhibition. J Pharmacol Exp Ther. 355(3):397–409.

Foo LC, Allen NJ, Bushong EA, Ventura PB, Chung WS, Zhou L, Cahoy JD, Daneman R, Zong H, Ellisman MH, Barres BA. 2011. Development of a method for the purification and culture of rodent astrocytes. Neuron 71(5):799–811.

Gilliam D, O’Brien DP, Coates JR, Johnson GS, Johnson GC, Mhlanga-Mutangadura T, Hansen L, Taylor JF, Schnabel RD. 2014. A homozygous KCNJ10 mutation in Jack Russell Terriers and related breeds with spinocerebellar ataxia with myokymia, seizures, or both. J Vet Intern Med 3:871–7.

Gomez RC, Wawro P, Lis P, Alessi DR, Pfeffer SR. 2019. Membrane association but not identity is required for LRRK2 activation and phosphorylation of Rab GTPases. J Cell Biol 218:4157–4170.

Gonzalez-Reyes LE, Verbitsky M, Blesa J, Jackson-Lewis V, Paredes D, Tillack K, Phani S, Kramer ER, Przedborski S, Kottmann AH. 2012. Sonic Hedgehog Maintains Cellular and Neurochemical Homeostasis in the Adult Nigrostriatal Circuit. Neuron 75:306–319.

Greggio E, Jain S, Kingsbury A, Bandopadhyay R, Lewis P, Kaganovich A, van der Brug MP, Beilina A, Blackinton J, Thomas KJ, Ahmad R, Miller DW, Kesavapany S, Singleton A, Lees A, Harvey RJ, Harvey K, Cookson MR. 2006. Kinase activity is required for the toxic effects of mutant LRRK2/dardarin. Neurobiol Dis 23:329–341.

Guo J, Otis JM, Higginbotham H, Monckton C, Cheng J, Asokan A, Mykytyn K, Caspary T, Stuber GD, Anton ES. 2017. Primary Cilia Signaling Shapes the Development of Interneuronal Connectivity. Dev Cell 42:286-300.e4.

Hill SA, Blaeser AS, Coley AA, Xie Y, Shepard KA, Harwell CC, Gao W-J, Garcia ADR. 2019. Sonic hedgehog signaling in astrocytes mediates cell type-specific synaptic organization. Elife 8:e45545.

Iannotta L, Biosa A, Kluss JH, et al. 2020. Divergent Effects of G2019S and R1441C LRRK2 Mutations on LRRK2 and Rab10 Phosphorylations in Mouse Tissues. Cells 9(11):2344.

Inyushin M, Kucheryavykh LY, Kucheryavykh YV, Nichols CG, Buono RJ, Ferraro TN, Skatchkov SN, Eaton MJ. 2010. Potassium channel activity and glutamate uptake are impaired in astrocytes of seizure-susceptible DBA/2 mice. Epilepsia 9:1707–13.

Ishii S, Sasaki T, Mohammad S, Hwang H, Tomy E, Somaa F, Ishibashi N, Okano H, Rakic P, Hashimoto-Torii K, Torii M. (2021). Primary cilia safeguard cortical neurons in neonatal mouse forebrain from environmental stress-induced dendritic degeneration. Proc Natl Acad Sci U S A 118:1.

Ito G, Katsemonova K, Tonelli F, Lis P, Baptista MAS, Shpiro N, Duddy G, Wilson S, Ho PW-L, Ho S-L, Reith AD, Alessi DR. 2016. Phos-tag analysis of Rab10 phosphorylation by LRRK2: a powerful assay for assessing kinase function and inhibitors. Biochem J 473:2671–2685.

Jaleel M, Nichols RJ, Deak M, Campbell DG, Gillardon F, Knebel A, Alessi DR. 2007. LRRK2 phosphorylates moesin at threonine-558: characterization of how Parkinson’s disease mutants affect kinase activity. Biochem J. 405:307–17.

Kalogeropulou AF, Freemantle JB, Lis P, Vides EG, Polinski NK, Alessi DR. 2020. Endogenous Rab29 does not impact basal or stimulated LRRK2 pathway activity. Biochem J. 477(22):4397–4423.

Kasahara K, Miyoshi K, Murakami S, Miyazaki I, Asanuma M. 2014. Visualization of astrocytic primary cilia in the mouse brain by immunofluorescent analysis using the cilia marker Arl13B. Acta Med Okayama 68:317–22.

Khakh BS, Deneen B. 2019. The Emerging Nature of Astrocyte Diversity. Annu Rev Neurosci 42:187–207.

Khan SS, Dhekne HS, Tonelli F, Pfeffer, SP. 2020. Analysis of Primary Cilia in Rodent Brain By Immunofluorescence Microscopy. protocols.io https://dx.doi.org/10.17504/protocols.io.bnwimfce

Kuwahara T, Inoue K, D’Agati VD, Fujimoto T, Eguchi T, Saha S, Wolozin B, Iwatsubo T, Abeliovich A. 2016. LRRK2 and RAB7L1 coordinately regulate axonal morphology and lysosome integrity in diverse cellular contexts. SciRep 6:29945

Lis P, Burel S, Steger M, Mann M, Brown F, Diez F, Tonelli F, Holton JL, Ho PW, Ho SL, Chou MY, Polinski NK, Martinez TN, Davies P, Alessi DR. 2018. Development of phospho-specific Rab protein antibodies to monitor in vivo activity of the LRRK2 Parkinson’s disease kinase. Biochem J. 475(1):1–22.

Ordónez LAJ, Fernández B, Fdez E, Romo-Lozano M, Madero-Pérez J, Lobbestael E, Baekelandt V, Aiastui A, López deMunaín A, Melrose HL, Civiero L, Hilfiker S. 2019. RAB8, RAB10 and RILPL1 contribute to both LRRK2 kinase–mediated centrosomal cohesion and ciliogenesis deficits. Hum Mol Genet 28:3552–3568.

Lee H, James WS, Cowley SA. 2017. LRRK2 in peripheral and central nervous system innate immunity: its link to Parkinson’s disease. Biochem Soc Trans 45:131–139.

Li X, Patel JC, Wang J, Avshalumov M V., Nicholson C, Buxbaum JD, Elder GA, Rice ME, Yue Z. 2010. Enhanced striatal dopamine transmission and motor performance with LRRK2 overexpression in mice is eliminated by familial Parkinson’s disease mutation G2019S. J Neurosci 30:1788–1797.

Linhart R, Wong S, Cao J, Tran M, Huynh A, Ardrey C, Park J, Hsu C, Taha S, Peterson R, Shea S, Kurian J, Venderova K. 2014. Vacuolar protein sorting 35 (Vps35) rescues locomotor deficits and shortened lifespan in Drosophila expressing a Parkinson’s disease mutant of Leucine-rich repeat kinase 2 (LRRK2). Mol Neurodegener 9:23.

Liu Z, Bryant N, Kumaran R, Beilina A, Abeliovich A, Cookson MR, West AB. 2018. LRRK2 phosphorylates membrane-bound Rabs and is activated by GTP-bound Rab7L1 to promote recruitment to the trans-Golgi network. Hum Mol Genet 27:385–395.

Madero-Pérez J, Fdez E, Fernández B, Lara Ordóñez AJ, Blanca Ramírez M, Gómez-Suaga P, Waschbüsch D, Lobbestael E, Baekelandt V, Nairn AC, Ruiz-Martínez J, Aiastui A, López de Munain A, Lis P, Comptdaer T, Taymans J-M, Chartier-Harlin M-C, Beilina A, Gonnelli A, Cookson MR, Greggio E, Hilfiker S. 2018. Parkinson disease-associated mutations in LRRK2 cause centrosomal defects via Rab8a phosphorylation. Mol Neurodegener 13:3.

Mandemakers W, Snellinx A, O’Neill MJ, de Strooper B. 2012. LRRK2 expression is enriched in the striosomal compartment of mouse striatum. Neurobiol Dis 48:582–593.

Martín R, Bajo-Grañeras R, Moratalla R, Perea G, Araque A. 2015. Circuit-specific signaling in astrocyte-neuron networks in basal ganglia pathways. Science 349:730–734.

Mir R, Tonelli F, Lis P, Macartney T, Polinski NK, Martinez TN, Chou MY, Howden AJM, König T, Hotzy C, Milenkovic I, Brücke T, Zimprich A, Sammler E, Alessi DR. 2018. The Parkinson’s disease VPS35[D620N] mutation enhances LRRK2-mediated Rab protein phosphorylation in mouse and human. Biochem J 475:1861–1883.

Nirujogi RS, Tonelli F, Taylor M, Lis P, Zimprich A, Sammler E, Alessi DR. 2021. Development of a multiplexed targeted mass spectrometry assay for LRRK2-phosphorylated Rabs and Ser910/Ser935 biomarker sites. Biochem J. 478(2):299–326.

Nwaobi SE, Cuddapah VA, Patterson KC, Randolph AC, Olsen ML. 2016. The role of glial-specific Kir4.1 in normal and pathological states of the CNS. Acta Neuropathol 132:1–21.

Pfeffer SR. 2018. LRRK2 and Rab GTPases. Biochem Soc Trans 46:1707–1712.

Pfeffer SR. 2017. Rab GTPases: master regulators that establish the secretory and endocytic pathways. Mol Biol Cell 28:712–715.

Poewe W, Seppi K, Tanner CM, Halliday GM, Brundin P, Volkmann J, Schrag A-E, Lang AE. 2017. Parkinson disease. Nat Rev Dis Prim 3:17013.

Purlyte E, Dhekne HS, Sarhan AR, Gomez R, Lis P, Wightman M, Martinez TN, Tonelli F, Pfeffer SR, Alessi DR. 2018. Rab29 activation of the Parkinson’s disease?associated LRRK2 kinase. EMBO J 37:1–18.

Rohatgi R, Milenkovic L, Scott MP. 2007. Patched1 Regulates Hedgehog Signaling at the Primary Cilium. Science 317:372–376.

Scholl UI, Choi M, Liu T, Ramaekers VT, Häusler MG, Grimmer J, Tobe SW, Farhi A, Nelson-Williams C, Lifton RP. 2009. Seizures, sensorineural deafness, ataxia, mental retardation, and electrolyte imbalance (SeSAME syndrome) caused by mutations in KCNJ10. Proc Natl Acad Sci U S A 106:5842– 584.

Sibille J, Pannasch U, Rouach N. 2014. Astroglial potassium clearance contributes to short-term plasticity of synaptically evoked currents at the tripartite synapse. J Physiol 592:87–102.

Sipos É, Komoly S, Ács P. 2018. Quantitative Comparison of Primary Cilia Marker Expression and Length in the Mouse Brain. J Mol Neurosci 64:397–409.

Sobu Y, Wawro PS, Dhekne HS, Pfeffer, SR. 2021. Pathogenic LRRK2 regulates ciliation probability upstream of Tau Tubulin kinase 2. Proc Natl Acad Sci. USA, in press. bioRxiv 2020.04.07.029983

Steger M, Diez F, Dhekne HS, Lis P, Nirujogi RS, Karayel O, Tonelli F, Martinez TN, Lorentzen E, Pfeffer SR, Alessi DR, Mann M. 2017. Systematic proteomic analysis of LRRK2-mediated rab GTPase phosphorylation establishes a connection to ciliogenesis. Elife 6:e31012.

Steger M, Tonelli F, Ito G, Davies P, Trost M, Vetter M, Wachter S, Lorentzen E, Duddy G, Wilson S, Baptista MAS, Fiske BK, Fell MJ, Morrow JA, Reith AD, Alessi DR, Mann M. 2016. Phosphoproteomics reveals that Parkinson’s disease kinase LRRK2 regulates a subset of Rab GTPases. Elife 5:e12813.

Sterpka A, Chen X. 2018. Neuronal and astrocytic primary cilia in the mature brain. Pharmacol Res 137:114–121.

Tong X, Ao Y, Faas GC, Nwaobi SE, Xu J, Haustein MD, Anderson MA, Mody I, Olsen ML, Sofroniew M V, Khakh BS. 2014. Astrocyte Kir4.1 ion channel deficits contribute to neuronal dysfunction in Huntington’s disease model mice. Nat Neurosci 17:694–703.

Waschbüsch D, Purlyte E, Pal P, McGrath E, Alessi DR, Khan AR. 2020. Structural Basis for Rab8a Recruitment of RILPL2 via LRRK2 Phosphorylation of Switch 2. Structure 28:406-417.e6.

West AB, Moore DJ, Biskup S, Bugayenko A, Smith WW, Ross CA, Dawson VL, Dawson TM. 2005. Parkinson’s disease-associated mutations in leucine-rich repeat kinase 2 augment kinase activity. Proc Natl Acad Sci USA 102:16842–16847.

West AB, Cowell RM, Daher JP, Moehle MS, Hinkle KM, Melrose HL, Standaert DG, Volpicelli-Daley LA. 2014. Differential LRRK2 expression in the cortex, striatum, and substantia nigra in transgenic and nontransgenic rodents. J Comp Neurol. 522(11):2465–80.

Zhai S, Tanimura A, Graves SM, Shen W, Surmeier DJ. 2018. Striatal synapses, circuits, and Parkinson’s disease. Curr Opin Neurobiol 48:9–16.

Zhang Y, Chen K, Sloan SA, Bennett ML, Scholze AR, O’Keeffe S, Phatnani HP, Guarnieri P, Caneda C, Ruderisch N, Deng S, Liddelow SA, Zhang C, Daneman R, Maniatis T, Barres BA, Wu JQ. 2014. An RNA-Sequencing Transcriptome and Splicing Database of Glia, Neurons, and Vascular Cells of the Cerebral Cortex. J Neurosci 34:11929–11947.

